# Microbial Production Of Ellagic Acid From Mango Pulp Processing Waste

**DOI:** 10.1101/2020.03.17.995597

**Authors:** Anandan Rubavathi, Athiappan Murugan, Kannan Visali

## Abstract

Ellagic acid has gained momentum recently due to its various properties like anti-mutagenic, anti-carcinogenic, anti-oxidant, and anti-viral and many other benefits to human health. The present study focused on the microbial production of ellagic acid from mango pulp processing industrial waste an alternate method for conventional chemical extraction. Our experiments demonstrated that the 100 μg/ml of ellagic acid was produced by *Micrococcus luteus* from 9% of mango pulp waste and the optimization of ellagic acid production with Pontecorvo medium supplemented with 5.0 g of ellagitannin has yielded 37.80 ± 0.30 mg/g at pH 5.0, temperature 30 °C, ammonium nitrate (nitrogen source), glucose (carbon source), with 1.5% of inoculums after 24 h of incubation. Ellagic acid synthesized was further confirmed with the standard ellagic acid. Applications like drought resistant in plants, anti-microbial activity, anti-parasitic activity and anti-cancer activities have been proven. Ellagic acid exhibited potential applications and further research in product development is promising.

## 1. Introduction

Ellagic acid (EA) is hydrolysable tannin present in the majority of the plants and synthesized enzymatically from ellagitannins (ETs) [1] (Figure 7). EAs have wide applications such as anti-oxidant, anti-tumoural, anti-viral, anti-microbial, anti-inflammatory, and anti-carcinogenic activities such as anti-hepatotoxic, anti-cholesteric, anti-fibrogenic and anti-hepatocarcinogenic properties [2]. EA acts as a remarkable de-pigmentation, gas protective agent and reduces birth defects. Ellagic acid reacts in the human body that may promote good health and effective in the prevention of cancer, heart diseases and other chronic diseases. Nowadays, the demand for natural ellagic acid is increasing due to their use in the functional food industries as well as in pharmaceutical industries. Ellagitannins and ellagic acid interact with the cell walls or sites to facilitate to form complex protein, which prevents the formation of tumours by preventing and proliferation of metastatic cells [3, 4].

It has been chemically synthesised for more than two decades. However, chemical synthesis involves harsh treatments that are highly aggressive and leads to corrosion of vessels [5]. EA is being extracted by solvent extraction with concentrated H_2_SO_4_ or HCl in the industrial scale [6]. Further, recovering EA directly from plants would increase production cost as well as loss of biodiversity [7]. Alternatively, tannin rich mango pulp processing industrial waste can be used as substrate for bio-conversion into ellagic acid. Hence, the present work is aimed at producing ellagic acid from ellagitannins by the microbial bio-conversion [8].

Agricultural residues are the main effective source of natural bioactive compounds [9]. Mango wastes are used by the microbes for their own metabolism and finally produce some bioactive compounds that are important for soil health, plant growth and overall eco-balance. The utilization of agriculture wastes as abundant, bio-renewable and low-cost sources for the production of high value-added products are still under investigation, with limited outcomes. Researches in this field of therapeutic molecules extracted from agro-industrial waste sound good due to their eco-friendly and improve economy of agri-business.

Many researchers have already been studied on potential sources of bioactive compounds from agricultural by-products. The food industry is facing a great challenge to handle large quantity of waste generated from product recovery. Billions of dollars are spent on the treatment of agricultural and food waste to recover from the risk caused by these wastes to the environment. Therefore, the risks can be reduced and the costs for treatment of waste can also be reduced. Value for agricultural and product development would be possible through this bioconversion. Mango peels, mango seeds and other wastes are discarded in the industrial process of pulp recovery [10].

Mango peel can be used as a valuable, economic and cheaper source for the commercial production of the ellagic acid [11]. Ellagic acid can be obtained by the enzymatic hydrolyzes of ellagitannins. Therefore, the aim of this study was to elucidate the bacterial bio-conversion of ellagitannins into ellagic acid by submerged fermentation. Earlier many reports had studied the role of ellagitannase on ellagitannins biodegradation. Microbial conversion of ellagic acid from pomegranate husk had shown 65.7 mg/g [8]. Therefore, the aim of this study was to elucidate the bacterial bio-conversion of mango ellagitannins into ellagic acid using submerged fermentation.

## 2. Materials and methods

### 2.1. Collection of samples

The mango pulp wastes were collected from Adhiyamaan Mango pulp processing industry, Krishnagiri District, Tamil Nadu. The samples were dried, meshed and used for microbial conversion of ellagic acid.

### 2.2. Isolation and screening of ellagitannase producer

The soil samples were serially diluted up to 10^−6^ dilutions. The diluted samples were spread plated on to nutrient agar (NA) plates supplemented with 2% of ellagitannin of mango pulp processing waste and was incubated at 37 °C for 24h. After incubation, the bacterial colonies that were able to produce hallo-zone around the colonies treated as an ellagitannase positive colonies. Ellagitannase producers were screened with the help of methyl red (0.01 M methyl red in 0.01 N HCl) as pH indicator and flooded on the plates were kept for 5–10 min. Ellagitannase producers that showed red hallow zone around the colony were recorded as positive.

### 2.3. Production of ellagic acid

The ellagic acid producers were screened based on the diameter of the zone formed on the minimal salt medium (MSM) supplemented with different concentrations (1, 3, 5, 7, 9 and 10%) of mango pulp processing waste as substrate. Ellagic acid production was done using submerged fermentation with 5% of inoculums, and this setup was maintained at 37 °C, pH 7.0 for 24 h.

### 2.4. Optimization and production of ellagic acid

Erlenmeyer flasks (250 ml) containing 100 ml of Pontecorvo medium with different concentration of mango pulp wastes (0.05, 0.07, 0.10, 0.13 and 0.15 g) powder, carbon (sucrose, fructose, glucose, maltose, and xylose) (0.5 g) and nitrogen (peptone, yeast extract, beef extract, ammonium chloride and ammonium nitrate) (0.5 g) source were used for EA production by submerged fermentation of mango pulp waste with the microbial culture. A set up containing no substrate, carbon and nitrogen source has been treated as control. The submerged fermentation was carried out at various temperatures 25, 30, 35, 40 and 45 °C, for various incubation periods like 12, 24, 36, 48 and 72 h under constant 200 rpm agitation with various pH 4.0, 5.0, 6.0, 7.0 and 8.0 with the inoculums sizes of 0.1, 0.5, 1.0, 1.5 and 2.0%. The optimized condition was used for the production of ellagic acid from mango pulp waste powder using submerged fermentation.

### 2.5. Extraction and characterization of ellagic acid

Fermented broth containing ellagic acid was transferred to centrifuge tubes, and ellagic acid was separated by centrifugation at 2000 rpm for 20 minutes. The supernatant was transferred to separating funnel, and the equal volume of ethyl acetate was added to the supernatant and mixed well for separation. The upper aqueous layer containing ellagic acid was collected in a beaker and allowed for evaporation of ethyl acetate. About 2 ml of methanol was added after evaporation and stored in a refrigerator for further experimental analysis using Fourier transform infrared spectroscopy (FTIR).

### 2.6. Applications of ellagic acid

#### 2.6.1. Stress tolerance in plant

A pot experiment was conducted to investigate the physiological changes in *Vigna radiata* due to the exogenous application NaCl and protective effect of EA. Seeds were treated separately by soaking them for 6 h in a solution containing different concentrations of EA (0, 50, 100 and 200 μg/ml). Pre-treated seeds were sown in pots. Different concentrations of salts (0, 60 and 120 mM) were sprayed on the plants after a week of seed germination and continued for 4 weeks. Plant materials were collected after completion of experimental duration, and assessed for various parameters such as shoot length (cm), root length (cm), shoot fresh weight (g), root fresh weight (g), as well as a physiological parameter like chlorophyll content was measured.

#### 2.6.2. Estimation of chlorophyll

Chlorophyll contents were determined according to the method [12]. The fresh 1g leaves were homogenized in 20–40 ml in 80% acetone in a mortar and pestle. The homogenate was centrifuged at 3000 rpm for 30 min. The supernatant was collected, and the total volume was measured. About 5 ml of the properly diluted extract was taken to measure the absorbance at 645 nm, 663 nm (acetone blank) wavelength using a UV–vis spectrophotometer.

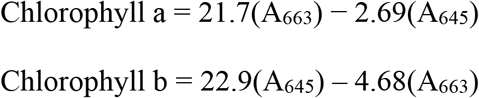

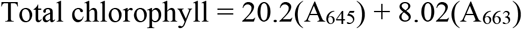

#### 2.6.3. Antimicrobial activity

The antibacterial activity of the ellagic acid was performed by the agar well diffusion method. Muller Hinton agar medium was inoculated with 0.1 ml of a fresh overnight culture of the *E. coli*, *Bacillus* sp., *Klebsiella pneumonia* and *Staphylococcus aureus* for testing the property of EA antibacterial test and the property of EA. Wells of 6 mm in diameter were punched in the agar and filled with different concentrations (100, 50, 25, 12.5 and 6.75 μg) of the EA. The plates were incubated at 37 °C for 24 h. Then, they were examined for the zone of inhibition, and the results were recorded. The anti-bacterial activity of ellagic acid was measured based on the diameter of the zone of inhibition.

#### 2.6.4. Anti parasitic properties-Larval toxicity test

A laboratory-reared larva was used for the evaluation of the larvicidal activity of ellagic acid. Twenty individuals of first, second, third and fourth instars larvae were kept in a 100 ml glass beaker containing 49 ml of de-chlorinated water and various concentrations of ellagic acid in different concentrations (50, 100 and 200 μg) were added. Larval food was given for the test larvae. The control was set up by 50 ml of de-chlorinated water alone. The control mortalities were corrected by using Abbott’s formula [13].

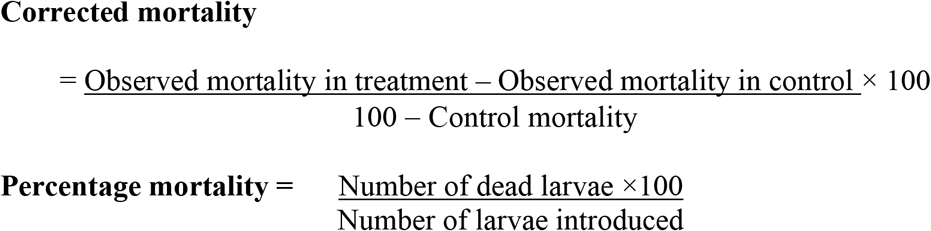

#### 2.6.5. Anti-cancer activity

The anti-cancer activity was analyzed with various cell lines (MCF-7, HT29, A549, MDA-MB-231 and PA1). The 3-(4,5-dimethylthiazol-2-yl)-2,5-dimethyltetrazolium bromide (MTT) assay, based on the conversion of yellow tetrazolium salt-MTT to purple-formazan crystals by metabolically active cells, provides a quantitative determination of viable cells (Morgan 1998). MCF-7, HT-29, A549, PA1 and MDA-MB-231 the human cancer cells obtained from the Korean cell line bank Seoul, Korea. Cells were plated onto 96-well plates at a cell density of 2 × 10^−5^ml^−1^ per well in 100 μl of RPMI 1640 and allowed to grow in a CO_2_ incubator for 24h (37 °C, 5% CO_2_). The medium then removed and replaced by fresh medium containing different concentration of sample for 24 h. The cells were incubated for 24–48 h (37 °C, 5% CO_2_). Then 20 μl MTT stock solutions (5 mg/ml in PBS) was added to each well and incubated for 5 h. The medium was removed and 200 μl DMSO added to each well to let dissolve MTT metabolic product. Then the plate shaken at 150 rpm for 5 min, and the optical density was measured at 620 and 570 nm. Untreated cells (basal) were used as a control of viability (100%), and the cell viability was calculated as follows:

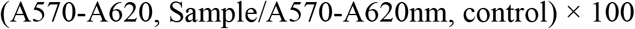

## 3. Results

### 3.1. Isolation and screening of ellagic acid producing bacteria

The soil samples were collected from 8 different site (Garden site GS1, GS2, Periyar University PU3, PU4, PU5, agriculture field AG6, AG7, AG8), and nearly 200 different colonies were isolated by standard plate count method. The number of colonies was counted for each sample at 10^−6^ dilutions. Nearly 15 bacterial isolates were shown positive for tannase production, of which one isolate PU4 was able to produce a 15 mm wider zone than other strains. Hence, the bacterial culture PU4 has been chosen for further experiment analysis.

### 3.2. Production of ellagic acid

Ellagic acid production was accomplished by inoculating MSM medium substituted with different concentrations of mango pulp processing waste (1%, 3%, 5%, 7%, 9%, and 10%) at 37 °C for 24 h. After fermentation, the cell growth was observed at 254 nm (OD) for ellagic acid production (Figure 2). The concentration of mango process was identified as 9% the production of ellagic acid taken for further scale-up production and extraction. It was calculated from the standard graph that 100 μg/ml of ellagic acid was produced when 9% of mango pulp was used as a substrate.

### 3.3. Optimization of ellagic acid

To optimize the media for maximum production of EA components such as different carbon and nitrogen sources were optimized. Even various temperatures, pH, incubation period and inoculums concentration were maintained. Among several carbon and nitrogen sources used for the EA production, glucose had yielded 24.68 mg/g and ammonium nitrate 24.52 mg/g of EA. Among various temperatures used for the EA production, 30 °C showed a maximum (24.38 mg/g) of EA production. Among various pH used for the EA production, pH 7.0 showed a maximum 24.51 mg/g, in case of incubation period at 24 h, the EA production is 24.37 mg/g and the size of the inoculums (1.5%) production of EA is 24.5 mg/g. The production of EA after optimization was about 37.80 ± 0.30 mg/g that means a 1.3-fold increase in production.

### 3.4. Characterization of ellagic acid

Characterization of the ellagic acid was done by using FTIR by their functional groups. The test compound was analyzed by FTIR, the results of the spectrogram represent the presence of glycosidic groups V _(O-H)_ that is the presence of OH stretching with a broad peak at 3554 cm^−1^. The carboxylic substituent’s C=O stretching peak at 1691.57 cm^−1^. The amide group at 1193– 1057 cm^−1^ and the alkaloids at 638.44 cm^−1^ correspond to ellagic acid. All these bands are more or less similar to the standard ellagic acid (Figure 1). Based on the functional group and the chemical structure of the extracted compound, it may be presumed as ellagic acid. The sample was further confirmed with the comparison with standard ellagic acid.

**Figure 1:**
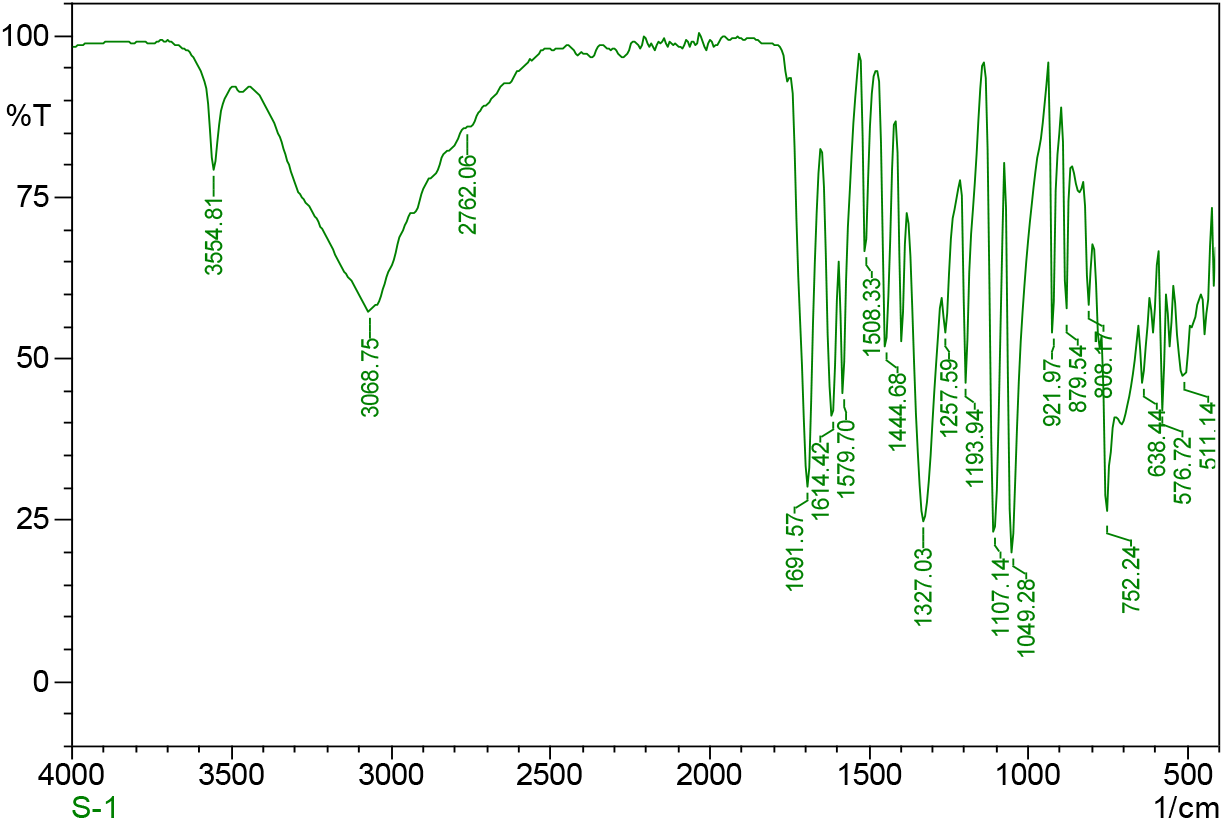
Characterization of ellagic acid produced by the natural isolates using FTIR

**Figure 2:**
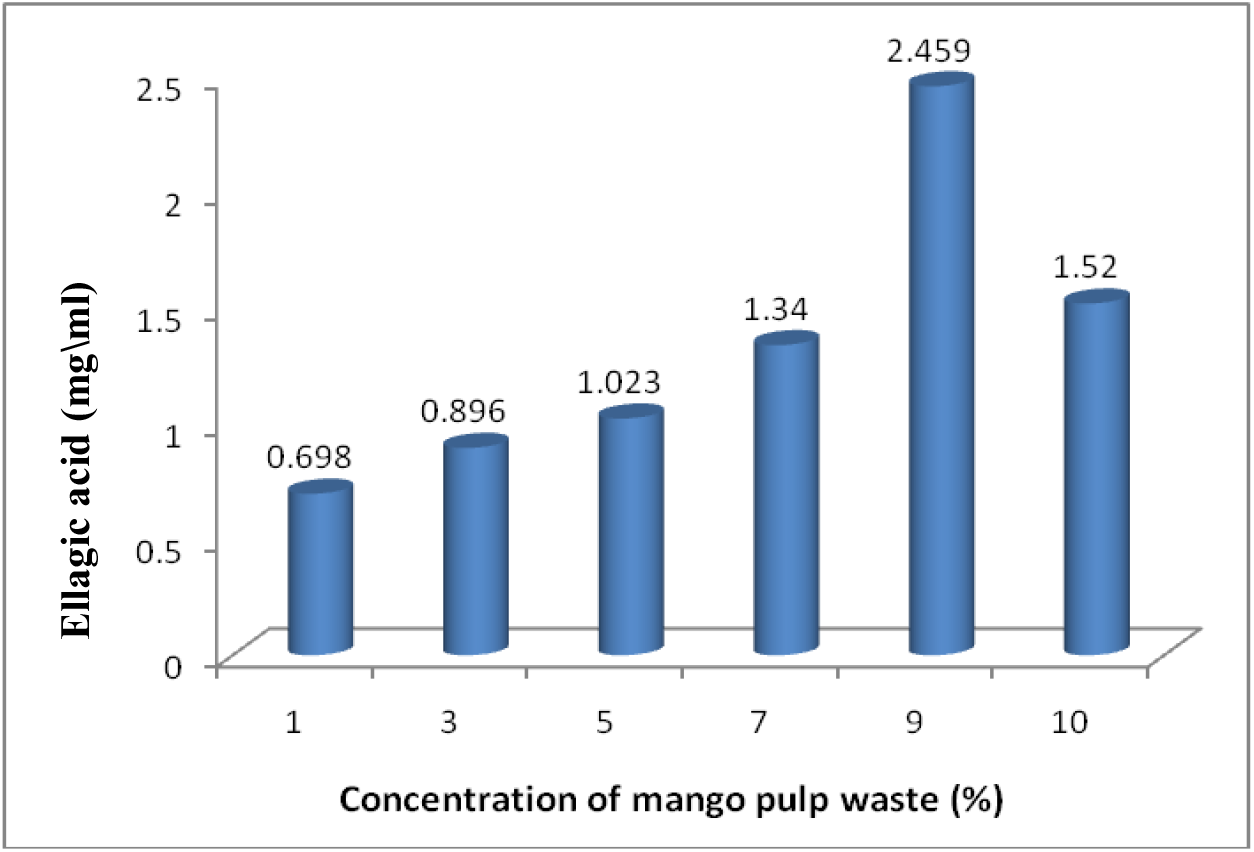
Effect of different concentration of mango pulp waste on the ellagic acid production

### 3.5. Applications of ellagic acid

#### 3.5.1. Stress tolerance in plant

Salt stress severely affects *Vigna radiata* (green gram) varieties although the effects of NaCl on the shoot and root length varied among green gram varieties (Figure 3). Salt stress had a diminished shoot fresh and dry weight as well as root fresh and dry weight. Treating seeds with EA had a pronounced effect on the above parameters, while the control plant was severely affected by salt treatment. The highest concentration of 200μg/ml EA reduced salinity effects of NaCl treatments in plants, resulting in longer shoots when compared to other concentrations of NaCl. Whereas medium concentrations of 55μg/ml EA had no effect in plants, and the concentration of NaCl was maintained. Root length was generally variable in all salt treatments and little influenced by the addition of EA in *Vigna radiata*.

**Figure 3:**
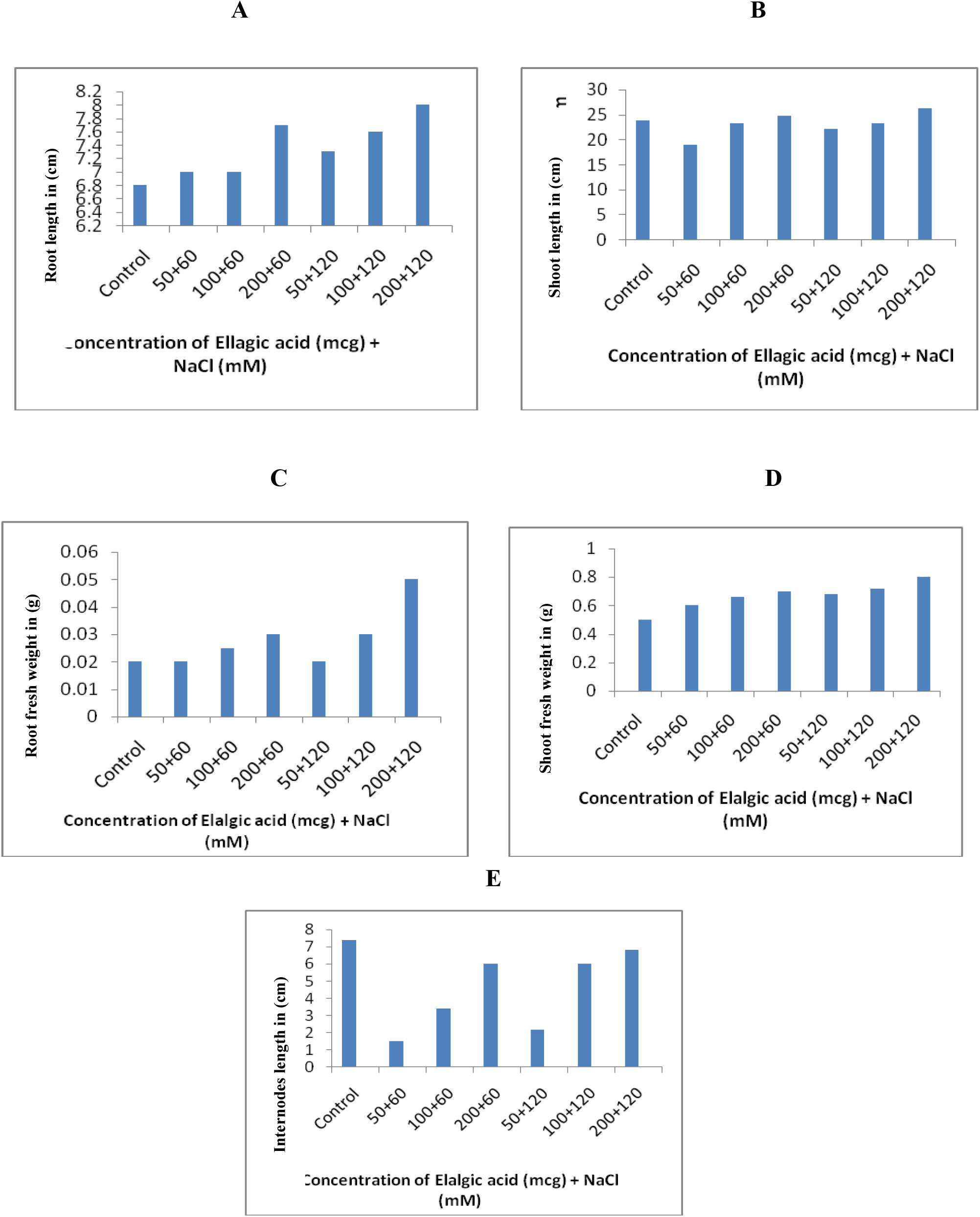
Drought resistant properties of the ellagic acid on green gram plants: root length (A) and shoot length (B), fresh root (C) and fresh shoot (D) and inter-node (E) *Vigna radiata* of under non-saline (control) and saline (60 mM and 120 mM NaCl) conditions.

#### 3.5.2. Chlorophyll content of plant

Application of EA evaluated the effect of salinity and enhanced the growth of the plants. Among different concentrations of EA, 200μg/ml had increased the chlorophyll content of leaf under saline conditions at *Vigna radiata* (green gram) at 120 mM of salinity compared with others. With regard to chlorophyll b, 200 μg/ml of EA increased the content in *Vigna radiata* (green gram) at 120 mM NaCl but not at 60 mM NaCl, while in *Vigna radiata* (green gram), 200 μg of EA was more effective at both salinity treatments. Salinity stress also reduced the total chlorophyll amounts and the application of EA alleviated the effect of salinity when applied at 200 μg/ml at *Vigna radiata* (green gram), an increase of the total chlorophyll was also observed after EA addition, particularly at high concentration while (Figure 4).

**Figure 4:**
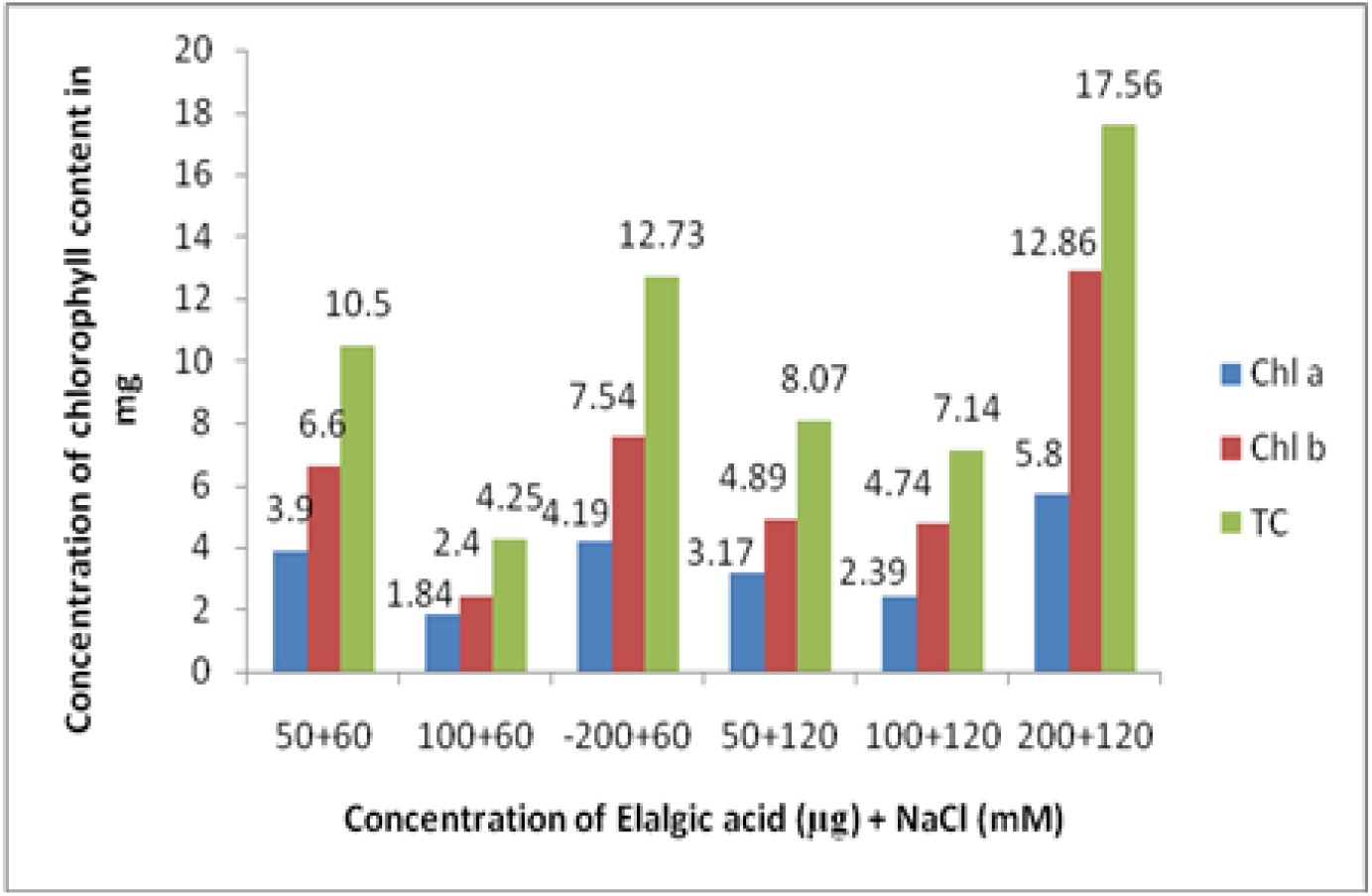
Effect of EA on chlorophyll a, chlorophyll b (b) total chlorophyll (TC) (c) of *Vigna radiata* (green gram) under saline and non-saline conditions.

#### 3.5.3. Anti-microbial activity

The antimicrobial activity of the ellagic acid was assessed by testing the organism for antimicrobial sensitivity, and the pathogen was found to be resistant to the drug used. The antimicrobial activity of the extract increased linearly with the increase in the concentration of extract (μg/ml). As compared with standard drugs, the result revealed that in the extract the bacterial activity for *Staphylococcus aureus* (23 mm) and *Bacillus* sp. (25 mm) was sensitive as compared to *E. coli* and *Klebsiella pneumonia* (Results not shown).

#### 3.5.4. Larvicidal activity

The larvicidal activity of ellagic acid was tested with mosquito larvae (*Aedes aegypti*). It was observed that 15% mortality was noted on the 2nd day after treatment with 50 μg of EA while it was 50% mortality with 200 μg EA. The mortality was 100% after 4th day and 5th-day treatment with 200 μg and 100 μg, respectively. In comparison to standard ellagic acid (100 μg), which gave 100% mortality on the third day, it was observed that ellagic acid produced from mango pulp waste gave 100% mortality on the 5th day when 100 μg was used (Figure 5).

**Figure 5:**
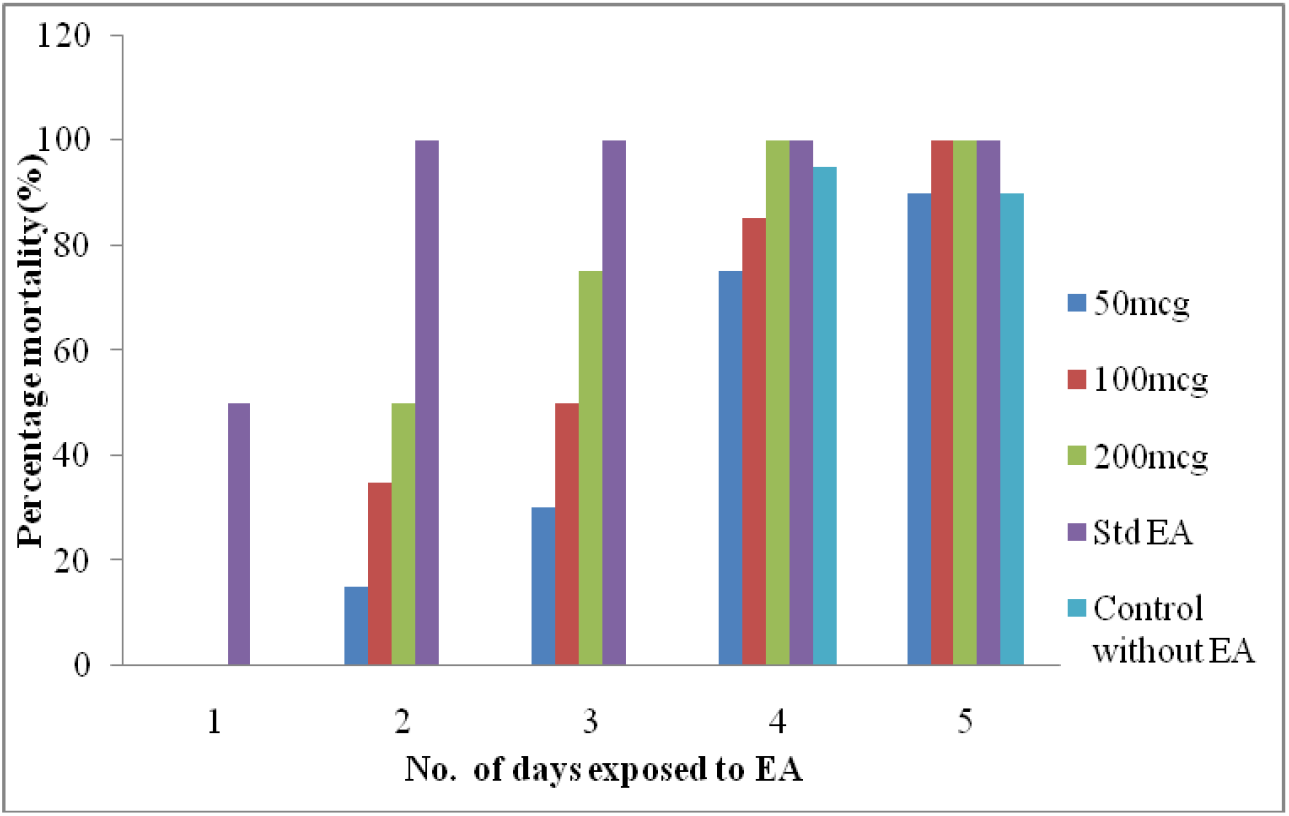
Larvicidal Activity of ellagic acid at different concentration test against *Aedes aegypti*

#### 3.5.5. Anti-cancer activity

Ellagic acid was screened using a cell viability assay (MTT assay) to examine the anti-cancer activity with various cell lines such as MCF-7-Breast cancer cell line, HT-29-Colon Cancer cell line, A549 – Lung Cancer cell line, PA-1 – Ovarian Cancer cell line, MDA-MB-231 – Breast Cancer cell line for 72 h. Cell lines were treated with different doses of ellagic acid and thymoquinone for 72 h, and then anticancer efficacy was observed for each cell line.

#### PA-1-Ovarian Cancer Cell line

The IC50 value for this cell line was 18.74μg/ml, which is between the therapeutic index of 17.82–19.71 μg/ml and this was compared with thymoquinone (16.11 μg/ml) (Figure 6).

**Figure 6:**
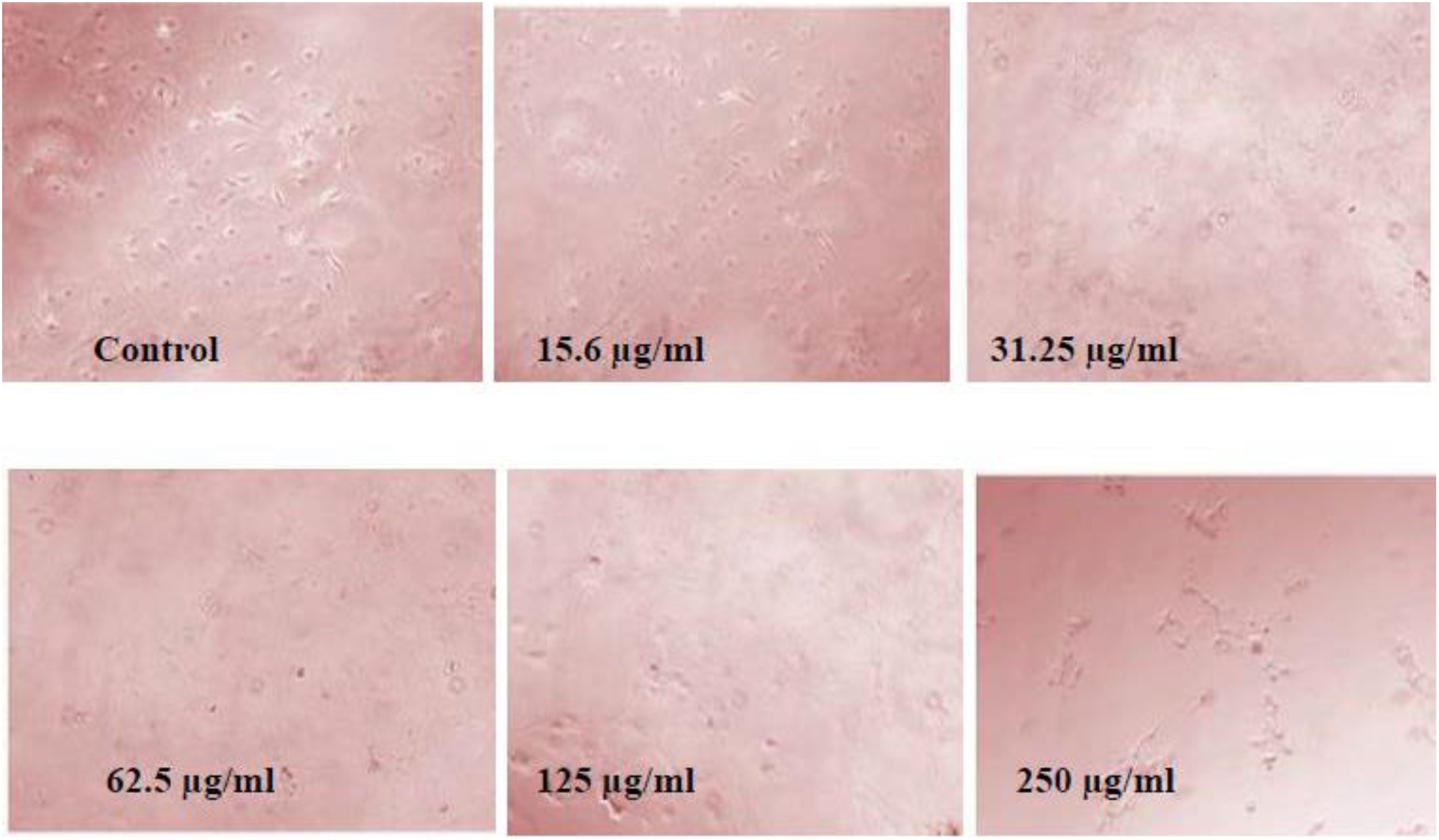
Anticancer activity of ellagic acid at the different concentration tested against PA1 cell lines

**Figure 7:**
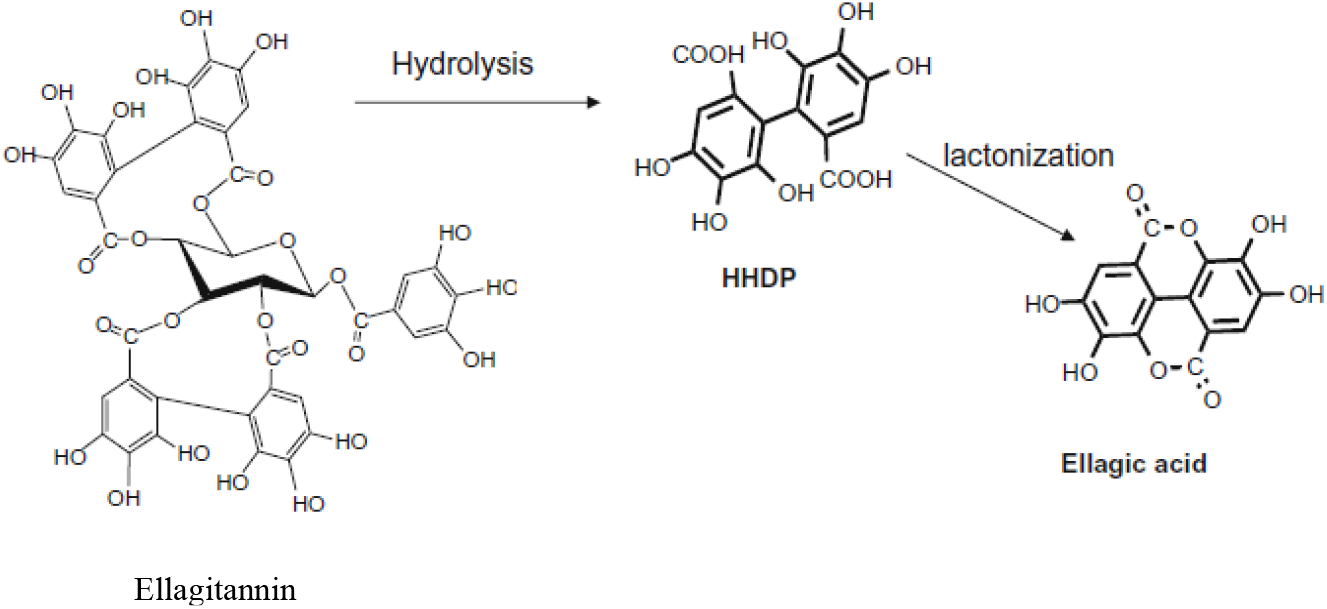
Schematic showing enzymatic hydrolysis of ellagitannin

## 4. Discussion

The bioactive compounds are rich in plants and used both in traditional and modern medicine against various ailments. However, bioactive compounds available in many plants are not fully utilized and remains as an unexplored area. Ellagic acid is a polyphenol, in particular, an ellagitannin, which can be found in a great proportion in many plants, has been known for its health benefits many years ago. Researchers have started work on pharmaceutical applications of ellagic acid only in recent years [14]. The phenolic nature of ellagic acid has the capacity to form complexes with other molecules and influence positive health benefits. Anticancer property against breast cancer, prostate cancer and colon cancer had been known for many years [15].

Bioavailability of ellagic acid has become as a great challenge, and it has become a hurdle for in vitro studies and in vivo studies [16]. There is an urgent need for the formulation of a novel method for the production of ellagic acid using microbial sources using the cheaper resource as a substrate. Ellagitannins of mango chosen for ellagic acid production based on earlier reports stated that strawberry, cranberry, blueberry and blackberry are a rich source of ellagitannin. Similarly, have attempted on the ellagic acid production from the aqueous extract obtained from pomegranate husk (*Punica granatum*) and from acron fringe oak [17]. Many similar other agro wastes can be used for the production of ellagic acid which is rich in ellagitannins and mango pulp processed waste was used for this study.

Similarly, about 65.7 mg/g of ellagic acid produced with 7 % of pomegranate husk was the substrate [18]. Pomegranate has more amount of ellagitannin than mango pulp. Yet another study has used *A. niger* for the production of ellagic acid was able to produce only 21.19 mg/g with 7.5 g of pomegranate husk [19]. The amount of ellagic acid produced was low due to the tannin content of mango pulp waste also less. The previous had achieved 23.1% of ellagic acid from 12g of Cresetoe bush at 36 h using *A. niger* GH1 strain [17, 20]. *A. niger* produces tannin acyl hydrolase which are able hydrolyzes ellagitannin-rich plant materials [18, 21]. Similar studies had been conducted on the tanning degrading abilities of fungal strains GH1 and PSH [22].

Previous studies also have achieved 0.256 mg/l of ellagic acid with a temperature 35 °C using mahua bark and *Aspergillus awamori*. Some researchers reported that the rising temperature would affect the solubility of the protein, so that enzyme activity increased. But higher temperatures would denature the enzymes thus decreased the activity [23]. By using different ranges of all parameters in response surface methodology (RSM), the optimized condition for the high production of ellagic acid. The maximum ellagic acid production was observed with the following parameters glucose (0.83), ammonium nitrate (0.83), temperature (38.3 °C), pH (6.6), the incubation period (48 h), potassium chloride (0.37) and inoculums size (1.83%).

Using the optimized conditions, 496.75 mg/g of ellagic acid production was achieved which was 1.4 times increase in the production. The previous study had shown that *A. niger* HT4 which degraded the ellagitannins present in the pomegranate had produced 350.21 mg/g of EA [19]. Optimization of EA production using ellagitannin of MPPW as the substrate has yielded 37.80 ± 0.03 mg/g EA at 37 °C temperature, pH, carbon, nitrogen source [24]. There has been 2.3-fold increased production of ellagic acid (24.68 mg/g) than normal when glucose was used at 10.0 mg/g concentration.

The previous study reported that the ellagic acid production was carried out from the ellagitannins of containing the amount of ellagic acid was evaluated by HPLC. HPLC analysis of ellagic acid was displayed at 5.490 min with 254 nm [25]. Prior to the characterization, recovery of ellagic acid was carried out using ethanol formic acid extraction [19]. The characterization of ellagic acid obtained is important for understanding their properties and applications. It was confirmed by the comparison of standard ellagic acid. Ellagic acid is a complex metabolite therefore all the characterization has to carried out to confirm it with the standard molecule.

Also previous study reported that among different concentrations of EA, 110 μg/ml of EA increased the chlorophyll a under saline conditions at V1 and in V2 at 120 mM of salinity, With regard to chlorophyll b, 55 μg/ml of EA increased the content in V1 at 60 mM NaCl but not at 120 mM NaCl, while in V2, 110 ppm of EA was more effective at both salinity treatments. With regard to chlorophyll b, 200 μg/ml of EA increased the content in *Vigna radiata* (green gram) at 120 mM NaCl but not at 60 mM NaCl, while in *Vigna radiata* (green gram), 200 μg of EA was more effective at both salinity treatments [26].

Salinity stress also reduced the total chlorophyll amounts and the application of EA alleviated the effect of salinity when applied at 200 μg/ml at *Vigna radiata* (green gram), an increase of the total chlorophyll. Hence, from the previous study when compared to this study it confers that the ellagic acid produced the mango pulp processed waste has a better activity for the plant growth.

Antibacterial activity of ellagic acid has been studied using *Staphyllococcus epidermatis*, *B. cereus*, *K. pneumonia*, and *S. typhi*. The results proven that ellagic acid produced from the mango waste is potential antimicrobial agent. Similar other study had proven that that the ellagic acid of pomegranate and punicalagins significantly inhibited the growth of pathogenic *E. coli*, and *Pseudomonas aeruginosa* [27]. EA exhibited significant antibacterial activity than gentamycin and streptomycin when they were tested with above bacterial strains [28].

Previous study already reported that anti-larvicidal activity bioassays were also carried out using ellagic acid derivative compounds isolated from the bark of *Elaeocarpus parvifolius*, which showed potent anti-parasitic properties. A similar study was conducted by these compounds was evaluated for the power to inhibit *Babesia gibsoni*. This proves that ellagic acid has the anti-parasitic impact from the previous studies and this study [29]. A similar study was carried out by the larvicidal activities of *Tephrosia egregia* extract and its major component de-hydrorotenone was studied. High inhibitory activity was found for de-hydrorotenone and fuel and ester extracts from roots and stems, respectively.

Among the tested extracts, the dissolving agent extract from stems showed potent larvicidal activity (LC50 12.88 ± 0.64), whereas low activity was found for de-hydrorotenone [30].

However, the anti-cancer activity showed that the IC50 value of the ellagic acid is 46.08μg/ml, which is in between the therapeutic index of 43.2254.05 μg/ml. So, the sample has better activity for lung cancer (A549). Thymoquinone has an IC50 value of 43.90 μg/ml. Already found that A549 has the IC50 value >60 μg/ml [31]. Previous study reported that the A 549 cell line was resistant to anticancer compounds. It was found that IC50 value for A549 and RAW264.9 cells treated with ellagic acid nanocomposite was 72.3 and 24.8 μg/ml respectively, [32]. EA is one such compound that has been tested as a chemotherapeutic, radio sensitizer, antioxidant, etc. EA studies are performed on numerous growth varieties in laboratories which exhibit a big anti-carcinogenic activity. In some studies, evaluation is also performed in mice models. EA and radiation might persuade to be an efficient antitumor drug in clinics for the treatment of cancer.

The maximum amount of ellagic acid produced from the mango waste supplemented fermentation medium using native bacterial isolate. The ellagic acid produced showed anti-larval, anti-cancer, anti-microbial and stress-tolerant properties are promising drug candidates for future use. The current research data can appeal to smaller industries. Potential molecule like ellagic acid has great impact on the field of medicine, pharmaceutical, agriculture and food industries. Further, research on the molecular mechanism of various applications of ellagic acid would improve their wider usage.

## Acknowledgment

Both the authors acknowledge the DST-FIST facilities (Ref No. SR/FST/LSI – 640/2015C).

## Notes

The authors declare no competing financial interest.

